# Untangling direct species associations from mediator species effects with graphical models

**DOI:** 10.1101/470161

**Authors:** Gordana C. Popovic, David I. Warton, Fiona J. Thomson, Francis K. C. Hui, Angela T. Moles

**Affiliations:** School of Mathematics and Statistics and the Evolution & the Ecology Research Centre, UNSW Sydney, NSW 2052, Australia; Landcare Research, Lincoln, 7640, New Zealand; Mathematical Sciences Institute, The Australian National University, Acton, ACT 2601, Australia; School of Biological, Earth and Environmental Sciences & the Evolution & the Ecology Research Centre, UNSW Sydney, NSW 2052, Australia

## Abstract

Ecologists often investigate co-occurrence patterns in multi-species data in order to gain insight into the ecological causes of observed co-occurrences. Apart from direct associations between two species, two species may co-occur because they both respond in similar ways to environmental variables, or due to the presence of other (mediator) species.

A wide variety of methods are now available for modelling how environmental filtering drives species distributions. In contrast, methods for studying other causes of co-occurence are much more limited. “Graphical” methods, which can be used to study how mediator species impact co-occurrence patterns, have recently been proposed for use in ecology. However, available methods are limited to presence/absence data and methods assuming multivariate normality, which is problematic when analysing abundances.

We propose Gaussian copula graphical models (GCGMs) for studying the effect of mediator species on co-occurence patterns. GCGMs are a flexible type of graphical model which naturally accommodates all data types – binary (presence/absence), counts, as well as ordinal data and biomass, in a unified framework. Simulations for count data demonstrate that GCGMs are better able to distinguish effects of mediator species from direct associations than using existing methods designed for multivariate normal data.

We apply GCGMs to counts of hunting spiders, in order to visualise associations between species. We then analyze abundance data of New Zealand native forest cover (on an ordinal scale) to show how GCGMs can be used analyze large and complex datasets. In these data, we were able to reproduce known species relationships as well as generate new ecological hypotheses about species associations.

## INTRODUCTION

Understanding how and when species co-exist is central to understanding large scale patterns in biodiversity; ecosystem services such as pollination and seed dispersal, and may help scientists to predict how communities might re-assemble in response to climate change. There are three important drivers of co-occurrence. Firstly two species may co-occur because they both respond to environmental variables. For example, they might both be more abundant in warmer climates (Hawkins et al., 2003). Secondly, the species may both respond to the presence or abundance of other species, for example, they may both be hunted by the same predator (Gilinsky, 1984). Thirdly, species might have a direct association, such as facilitation, seed dispersal, or pollination (Freestone, 2006).

Methods for quantifying the effect of environmental variables on species’ co-occurrence are relatively well-established. The impact of measured environmental variables can be investigated using null models (Gotelli & Ulrich, 2010; Strong Jr et al., 2014; D’Amen et al., 2017), or using multivariate extensions of generalized linear models (GLMs), like in Pollock et al. (2014) and Hui et al. (2013). The impact of unmeasured environmental variables can be inferred using latent variable models (Warton et al., 2015; Ovaskainen et al., 2017). However, methods for estimating the effects of species on each others’ occurrence are much less well-developed.

To determine whether co-occurrence patterns between a pair of species are a result of relationships with mediator species, we can examine *conditional dependence* relationships between all the species. Conditional dependence describes how pairs of species are related, after controlling for all the other species in the dataset. Such relationships have not been widely investigated in ecology. Most of the research has instead focused on unconditional relationships based on correlations, including all the methods listed above. Markov models for binary data (Harris, 2016; Clark et al., 2018) have recently been proposed to uncover conditional relationships between species based on presence/absence data, although there is currently no framework to extend these to other data types. Additionally, for count data, Morueta-Holme et al. (2016) implemented a Gaussian graphical model on log transformed abundances. However, transforming count data can have undesirable effects on analyses (O’Hara & Kotze, 2010; Warton et al., 2016). This is especially the case for multi-species data (Warton & Hui, 2017) because counts are often small or zero, with not every species found at every sampling occasion. What is needed are tools for modelling conditional dependence capable of handling a broader range of distribution types, such that they can be directly applied to multi-species data without the need for data transformation.

In this paper we will demonstrate the use of Gaussian copula graphical models (GCGMs; Popovic et al., 2018) to uncover conditional relationships among species, and hence untangle the impact of mediator species on the co-occurrence patterns between pairs of species. Copula graphical models can naturally accommodate a wide variety of data types in a unified framework, including: binary variables (presence/absence); overdispersed counts; ordinal data; and biomass data. They do this by extending GLMs to accommodate multivariate data, essentially, by mapping GLM residuals onto the multivariate normal distribution. Using simulations, we show that GCGMs preform better at uncovering conditional relationships than log transformations of count data. We then demonstrate this method on a count dataset of spider abundances, and a dataset of ordinal cover categories of plants in New Zealand native forests. The latter dataset is particularly challenging due to its size (1311 taxa recorded at 964 sites) as well as the ordinal nature of the response; neither feature can be accommodated in any graphical modeling method currently used in ecology.

## MATERIALS AND METHODS

### Graphical models and co-occurrence

The concept of conditional dependence will be familiar to most applied researchers who have fitted linear models, where the term is typically used to refer to *controlling for* variables using a covariate. To measure if there is, for example, an effect of temperature on leaf width, after controlling for altitude, we can simply put both temperature and altitude in a model predicting leaf width, and test for an effect of temperature. This is the same to asking whether leaf width is conditionally dependent on temperature given altitude.

A similar strategy can be employed to uncover conditional dependence relationships between species, by using the species of interest as the response, and all the other species as predictors. This is in fact how Gaussian graphical models were first conceived and fitted (see: Banerjee et al., 2006; Meinshausen & Bühlmann, 2006). Modern methods for Gaussian graphical modelling use a LASSO-penalized likelihood approach (Friedman et al., 2008; Hastie et al., 2015) to estimate these conditional dependence relationships in Gaussian data, as they have better power and interpretablity, but the intuition remains the same.

Figure 1 demonstrates how graphical models can be used to disentangle whether correlations between species are due to relationships with mediator species or direct relationships between the species of interest. Here we are interested in the relationship between two species A and B, which we observe to be negatively correlated (red) in both scenarios given by the top and bottom rows. After controlling for a third species C (middle and right), we either find A and B are still related (top), or that their relationship was completely explained by both species responding to the mediator species C (bottom). It is important to emphasise that the only way to distinguish between these scenarios is to look at conditional relationships as given by the middle and right columns of Figure 1. The “graph” on the right, after which graphical models are named, is nothing but a convenient and visually appealing way to display these conditional dependence relationships. Conditional relationships are inferred from the matrix of partial correlations, which for Gaussian data is given by the inverse of the correlation matrix.

Even sophisticated methods for investigating species co-occurrence, like latent variable models, model dependence rather than conditional dependence relationships, and can therefore not untangle responses to mediator species from direct relationships between species. While graphical models are very rarely used in ecology, they are increasingly used in many other disciplines, from modelling gene networks (Krämer et al., 2009) to brain connectivity (Huang et al., 2010) and traffic flows (Sun et al., 2012).

**Figure 1:**
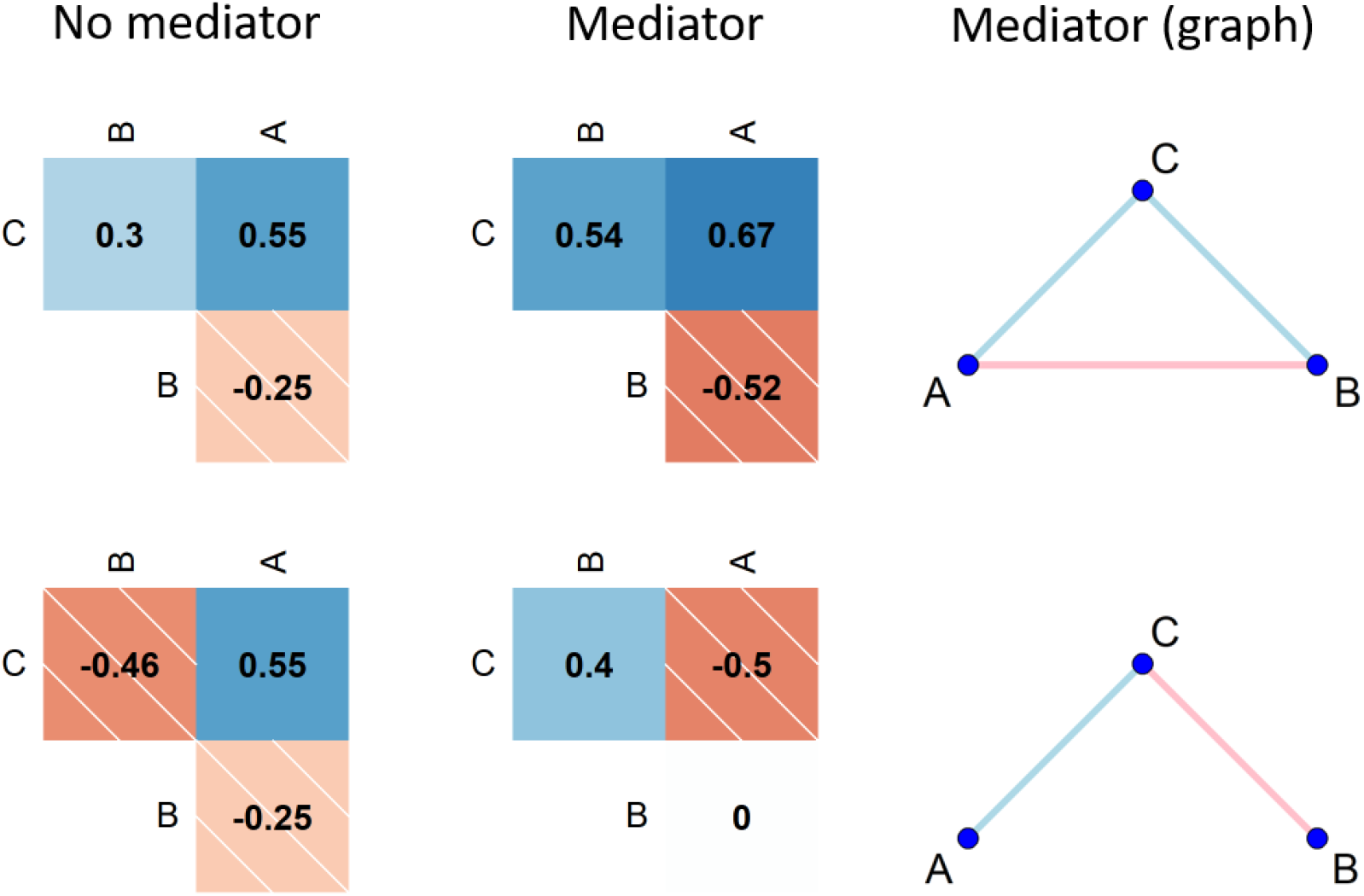
Left: correlations between species A, B, and C (red dashed for negative and blue for positive). Middle and right: partial correlations between each pair of species, after controlling for the other species. The middle column plots partial correlations in a matrix, while the third column plots this same information in a “graph”. Correlations (left) between species A and B are the same (*ρAB* = −0.25) in both scenarios (top and bottom). To see if this relationship is explained by both A and B responding to C, a mediator species, we can examine the conditional relationships (middle and right). In the bottom row, the relationship between A and B is fully explained by them both responding to C, while in the top row, A and B are related, even after controlling for C.

### Gaussian copula graphical models

Multi-species data (also known as multivariate abundance data) consist of measurements of abundance of plants or animals (most commonly presence/absence, counts, cover or biomass) simultaneously collected for a large number of taxa, classified to species or another taxonomic level. Gaussian graphical models cannot be directly used to model multi-species data, as the responses are almost always discrete. Gaussian copula graphical models provide a way to apply graphical models to any random variable, given a model for the statistical distribution of each species (its *marginal distribution*).

Copula models are so named because they couple a multivariate distribution (e.g. multivariate Gaussian) with any set of marginal distributions. For abundance data in ecology, which are generally discrete, the most useful marginal distributions are the binomial for presence/absence, the Poisson, negative binomial or binomial for counts, and a multinomial (often using a cumulative link model) for ordinal data. Copulas allow us to combine these into one model with the correlation structure of a multivariate Gaussian, which in turn allows us to indirectly apply Gaussian graphical models with all these data types. The term “Gaussian” in Gaussian copula graphical models refers only to the copula part of the model, the responses are modelled by the appropriate discrete distribution (*e.g*. negative binomial for counts). The Gaussian copula will be used here in preference to other copulas (*e.g*. vine copulas or archimedean copulas) as it allows the application of Gaussian graphical modelling algorithms in the estimation of the GCGMs (Popovic et al., 2018). Details of the model-fitting algorithm can be found in the Supplementary material, but briefly, it uses a Monte Carlo EM algorithm (Wei & Tanner, 1990), where repeated sets of Dunn-Smyth residuals (Dunn & Smyth, 1996) from the marginal model are used to estimate the copula likelihood, via importance sampling, with importance sampling weights updated iteratively following each graphical model fit to (re)weighted data, until convergence.

The GCGM model can easily incorporate any set of marginal distributions, including any GLMs, with an without covariates, and more complex models like generalised additive models, and cumulative link models for ordinal data. When using these models, we gain access to the tools of residual analysis and model selection to choose the appropriate marginal model for each species. This flexibility is the main advantage of GCGM over graphical modelling techniques previously used for multi-species data. The method implemented in Harris (2015) for example can only be used on presence/absence data, while the method of Morueta-Holme et al. (2016) requires multivariate normal data, or data that can be transformed to approximately satisfy multivariate normality.

With variables representing different species, one interpretation of the graphical model is as a model of species interactions; an attempt to identify which species interact directly with one another, and which are correlated due to their interaction with mediator species (Figure 3, row one v.s. row two). Other graphical methods used in ecology also make this connection (Harris, 2015; Morueta-Holme et al., 2016), however some care has to be taken with this interpretation (Dormann et al., 1028). Species which appear to interact may both be responding to unmeasured environmental variables. In addition, when species appear not to be interacting (partial correlation is zero), this should not be interpreted as evidence of no interaction, but rather that we do not have evidence of an interaction, perhaps because we do not have sufficient data to estimate these interactions (*e.g*. rare species had fewer connections in Figure 5, probably for this reason).

With these caveats in mind, we recommend GCGMs as a method for visualizing multivariate data, to be used in combination with other visualization methods like ordination. While species interactions cannot be directly inferred from observational data, GCGMs can be used as an exploratory tool to generate hypotheses about relationships between species, to inform further research.

### Data

We use two datasets in our analysis and simulations. The first is a dataset consisting of counts of hunting spiders caught in pitfall traps, with 12 species found at 28 sites (van der Aart & Smeenk-Enserink, 1974). The primary aim of this study was to identify the main environmental factors associated with the distribution of the species studied. The hunting spider data is a well known ecological dataset popularised in an important methodological paper (ter Braak, 1986). The data contain six covariates thought to be associated with spider abundance, namely: dry soil mass; percent cover of bare sand; percent cover of fallen leaves or twigs; percent cover of moss; percent cover of herb layer and reflection of the soil surface with a cloudless sky.

The second dataset consists of forest cover measurements collected at 964 native forest sites that form part of a network of permanent 20 x 20 m plots spread throughout New Zealand (Manaaki Whenua Landcare Research, 2018). A total of 1311 plant species were present in these plots, with the most common being herbs, graminoids, ferns, shrubs and trees. Plant cover (in ordinal categories) was assessed for each species in several tiers at different heights. The cover data we analysed were the maximum cover recorded over all the tiers. We use a subset of these data, which contains measurements at 964 sites identified as native forests. This reduced the number of species with at least one presence to 964.

## RESULTS

### Gaussian graphical model on transformed counts *vs*. GCGMs

We designed a simulation to compare the performance of a GCGM with appropriate count distributions for the responses (negative binomial) to the method in Morueta-Holme et al. (2016), where a Gaussian graphical model is used directly on log transformed counts. We know that transformations of this kind do not appropriately take the mean variance relationship of discrete data into account (O’Hara & Kotze, 2010), and would like to know what the implications of this are when studying conditional dependence relationships in count data. We explored this by simulating data in two groups of sites, with each species having a different mean across groups, and studied how well the two graphical modelling approaches recovered the true underlying graph from which the data were simulated.

We based the simulation on the hunting spiders abundance data (described in Methods). The precision matrix used for the simulation was obtained by fitting a GCGM to the data, This gives precision matrix is that approximately 50% sparse *i.e*., each species is conditionally dependent on about half the others. Sample sizes of 40, 100 and 400 were generated from this model. We generated data two groups, with different means for each species across groups of sites. Mean species abundances for one group were taken from the sample mean abundance in these data (and so differed by species). The species mean for the second group is the multiple of the first group mean by an effect size. We fitted a GCGM, and also a Gaussian graphical model fitted to residuals from linear regressions on (log + 1) transformed counts for each species. Performances of these methods was compared in terms of the number of correctly identified zero and non-zero conditional dependences (graph recovery rate).

As effect size increases, log transformation loses efficiency relative to the GCGM, as it does not adequately deal with the mean-variance relationship induced by the covariate effect (Figure 2). When effect size was one (*i.e*. there was no difference between groups), the two methods worked equally well. This is presumably because if there are no differences in means across sites, the mean-variance relationship implies no differences in variance either, so the equal-variance assumption of the Gaussian graphical model was satisfied when effect size was zero.

By better exploiting the structure of discrete data, using standard models for counts, GCGMs were more efficient at recovering the conditional dependence structure among variables than transforming counts to (approximate) normality.

**Figure 2:**
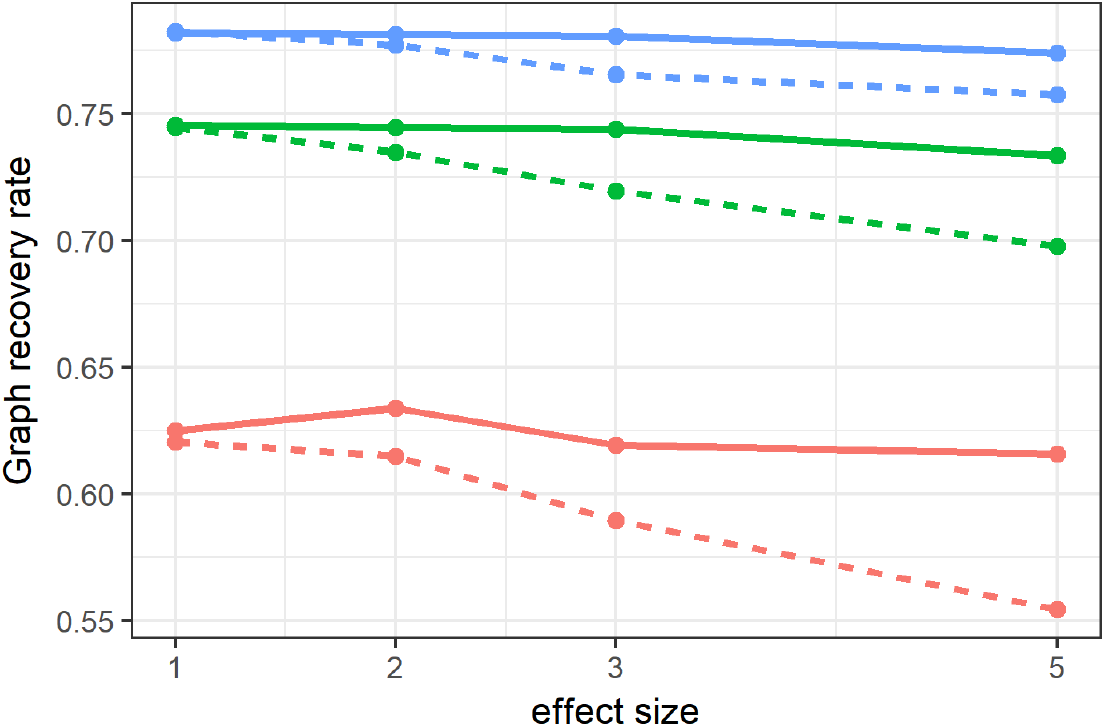
Graph recovery rates from simulations at different sample sizes (400 - blue, 100 - green, 25 - red) for graphical models constructed using (log+1) transformed counts (- - -) and a GCGM (—). Note that recovery rates are similar when there are no covariates (effect size=1), but that as effect size increases, the method using (log+1) transformed counts loses efficiency relative to the GCGM.

### GCGM analysis of counts

The first dataset we analyse are counts of hunting spiders (described in Methods). For these data we used negative binomial marginal distributions, as these can account for the overdispersion often observed in abundance count data. We can include predictors in these marginal distributions, and so will consider partial correlation both before and after accounting for observed environmental variables.

After controlling for the effect of other species (middle column Figure 3), partial correlations between species tend to be smaller, with many pairs of species being estimated to be conditionally independent. For these species, their correlation is explained by relationships with other species in the data. In addition, environment can explain a lot of (partial) correlation between species (bottom row have more zeroes than top row).

**Figure 3:**
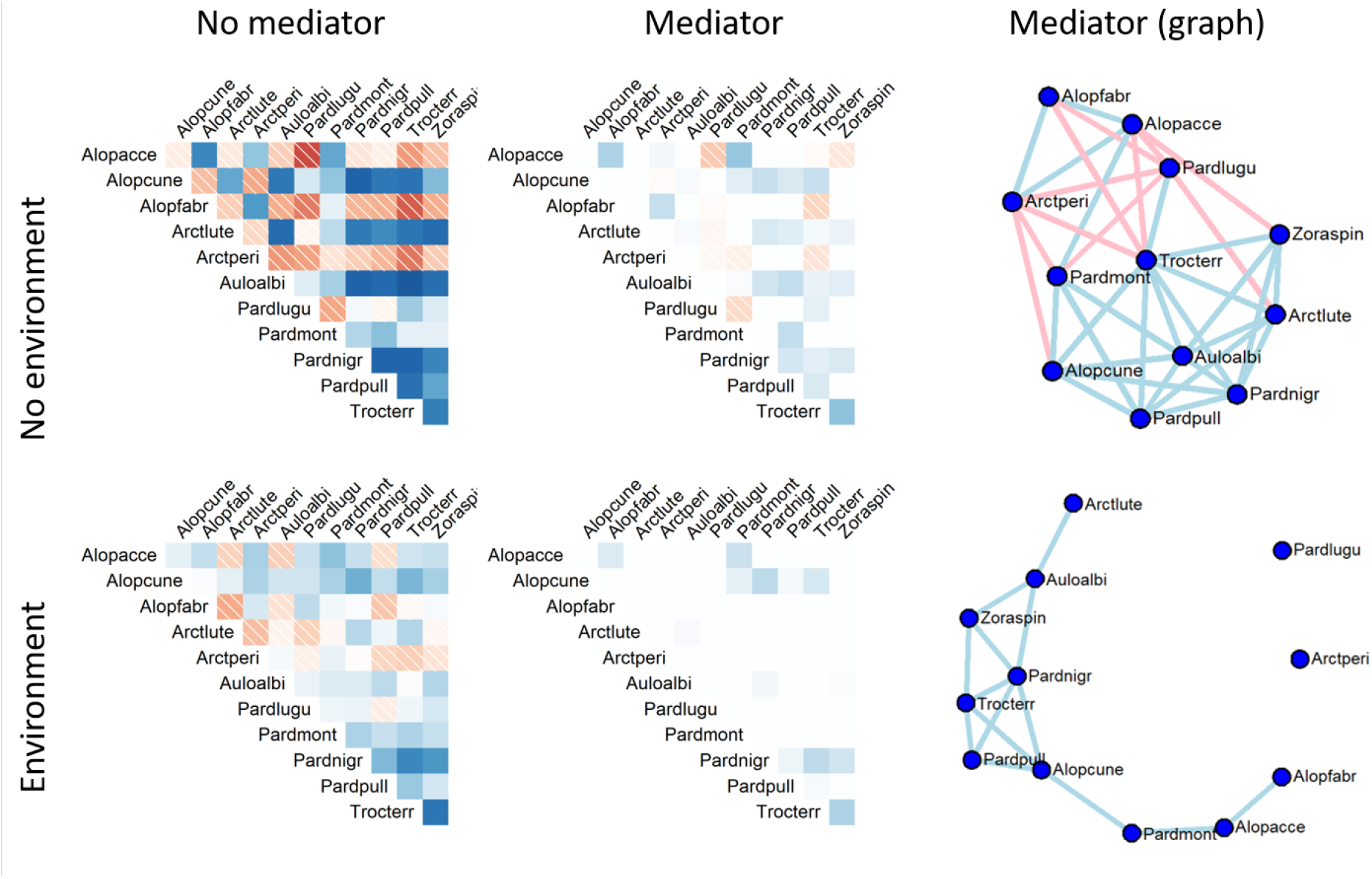
Raw and partial correlation between spider species. Correlations are positive (blue) if species co-occur more often than expected by their prevalence, negative (red, dashed) if they co-occur less often, and zero (white) if they co-occur at the rate expected by their prevalence. Correlations are plotted for the raw abundances (top) and after controlling for measured environmental variables (bottom). The left column shows raw correlations. Middle and right columns both display partial correlations, after additionally controlling for mediator species, with “graphs” on the right, which are plots of the network of conditional dependence. Names are abbreviations based on the first four letters of the genus, then the first for letters of the species names (see Supplementary material Appendix B for details).

Interpreting the graph of spiders, we observe that *Pardosa lugubris* (Pardlugu) has negative associations with both *Alopecosa accentuata* (Alopacce) and *Pardosa monticola* (Pardmont), who are positively associated, before controlling for covariates (Figure 3 top graph). However, these negative associations are absent after controlling for covariates (Figure 3 bottom graph). Looking at Figure 4, we can see that Pardlugu has decreased abundance in the presence of bare soil, while both Alopacce and Pardmont have the opposite response. The differences observed in the top and bottom of Figure 3 may therefore be due to these associations being explained by covariates.

**Figure 4:**
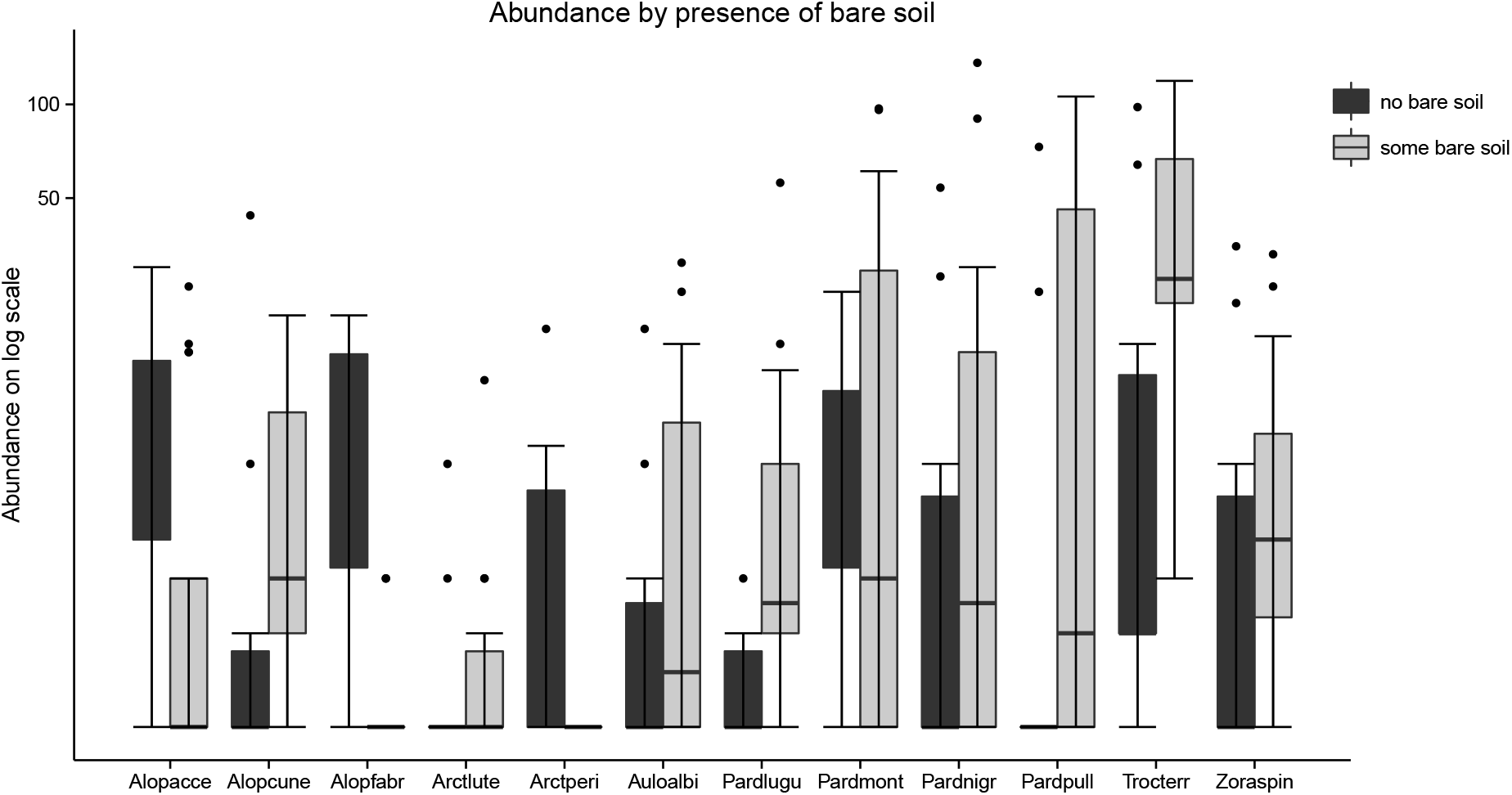
Spider abundance plotted against the presence of any bare soil. Notice that many of the species abundances differ according to the presence of any bare soil. For example, *Pardosa lugubris* (Pardlugu) has decreased abundance in the presence of bare soil, while both *Alopecosa accentuata* (Alopacce) and *Pardosa monticola* (Pardmont) have increased abundance.

### GCGM analysis of large ordinal dataset

We applied GCGMs to the New Zealand forest cover data (see Methods for description), a complex dataset that could not have been analysed using pre-existing methods.

In order to investigate associations between species in New Zealand native forests, we fitted a GCGM with cumulative link (Agresti, 2010) marginal distributions to these data. We included slope and altitude as covariates. The final graph was chosen by BIC.

Of the 964 species in the data, 142 were found to have non-zero partial correlations with other species. Due to the large number of species, the graph of all correlated species (Figure 5) is fairly complex, nevertheless we can see some important patterns. Most New Zealand forest species are positively associated with one another (blue lines), with the two ends of this gradient of positive association being negatively associated (as shown by the u-shaped group of positive associations dominating). The dominant species at one end of the spectrum tend to be associated with silver beech forest (*Lophozonia menziesii* (NOTMEN), *Raukaua simplex* (RAUSIM), *Myrsine divaricata* (MYRDIV), *Blechnum procerum* (BLEPRO), *Coprosma foetidissima* (COPFOE), and *Notogrammitis billardierei* (GRABIL)), while the other end of the gradient is dominated by species associated with the early-mid stages of regeneration of disturbed lowland forest (*Cyathea dealbata* (CYADEA), *Melicytus ramiflorus* (MELRAM), *Knightia excelsa* (KNIEXC) and *Uncinia uncinata* (UNCUNC)). This is partially consistent with our initial expectation that there would be a separation between the two main forest types of New Zealand. This prediction was based on the observation that podocarp-broadleaf forest and beech forests tend not to co-occur, but are not obviously separated by geography or climate.

**Figure 5:**
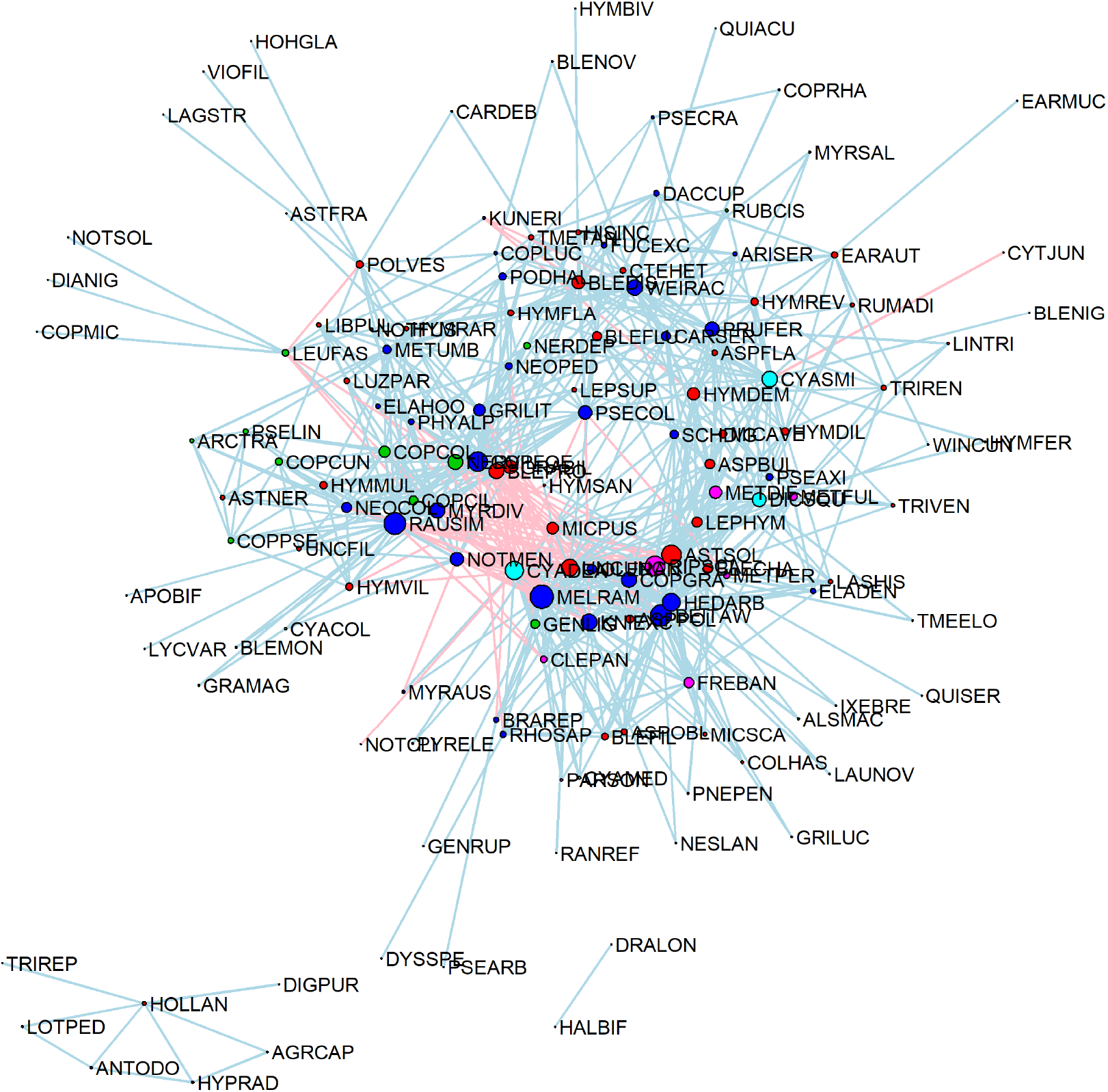
Graph for all species after controlling for covariates (slope and altitude) and mediating species. There are a range of positive (blue) and negative (red) partial correlations between species. The colour of the dot corresponds togrowth form; herbs (red), shrubs (green), trees (blue), vines (magenta) and tree ferns (cyan). To simplify the display for all plots, we excluded species that were conditionally independent from all other species. We employ the Fruchterman Reingold algorithm (Fruchterman & Reingold, 1991) to position nodes on the graphs in two dimensions. Some species codes reflect outdated taxonomy (e.g. Nothofagus menziesii has been renamed Lophozonia menziesii, but retains the NOT-MEN label in the dataset). Species at one end of the spectrum (NOTMEN, RAUSIM, MYRDIV, BLEPRO, COPFOE, and GRABIL) are associated with silver beech forest, while the other end of the gradient is dominated by species associated with the early-mid stages of regeneration of disturbed lowland forest (CYADEA, MELRAM, KNIEXC, and UNCUNC).

Some of the relationships we found were surprising, and can be used to generate hypotheses for further investigation. In Figure 5, the separation between the two main forest types of New Zealand was not supported for the three species in *Fuscospora*, the other genus of southern beech present in New Zealand *(Fuscospora solandri* (NOTSOL), *F. cliffortioides* (NOTCLI), *F. fusca* (NOTFUS). The three *Fuscospora* species fell in different parts of the graph rather than clustering together or with *Lophozonia menziesii*, and were not associated with many other species (either negatively or positively).

For more informative plots, we can “zoom in” on certain subsets of species, to better examine relationships between individual species. These subset graphs can display partial correlation between the species after accounting for covariates and all mediator species (including those not in the subset displayed).

We are able to distinguish between exotic herb species (*Trillium repens* (TRIREP), *Digitalis purpurea* (DIGPUR), *Holcus lanatus* (HOLLAN), *Agrostis capillaris* (AGRCAP), *Lotus pedunculatus* (LOTPED), *Anthoxanthum odoratum* (ANTODO), *Hypochaeris radicata* (HYPRAD) and *Libertia pulchella*(LIBPUL)), which are mostly positively correlated with one another, and negatively correlated with the native species (green text) by simply looking at the plot of raw correlations (Figure 6 left). This pattern is consistent with the fact that these exotic species are not shade tolerant, and thus differ from many of the native herbaceous species (a group which includes a range of understorey ferns) in not being able to survive in forest understoreys.

**Figure 6:**
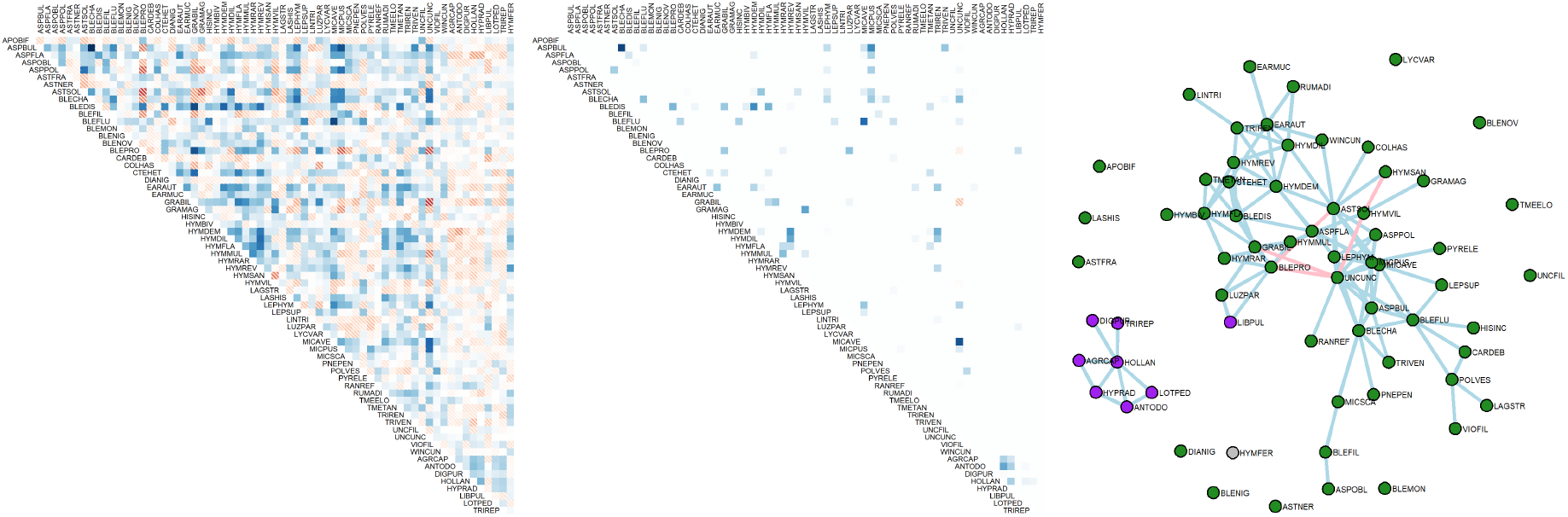
Herbs: partial correlations (middle and right) and correlations (left). The graph clearly shows a group of exotic (purple) herbaceous *(Trillium repens* (TRIREP), *Digitalis purpurea* (DIGPUR), *Holcus lanatus* (HOLLAN), *Agrostis capillaris* (AGRCAP), *Lotus pedunculatus* (LOTPED), *Anthoxanthum odoratum* (ANTODO) and *Hypochaeris radicata* (HYPRAD)) that are positively associated with one another, and not associated with any native (green) species. While all the plots reveal that the exotic species are distinct from the native species (rightmost eight species on left and middle plot), partial correlations additionally show that one exotic species *Libertia pulchella* (LIBPUL) is more closely related to the native species, than the other exotic species.

In addition to discerning the same group of exotic herbaceous species (purple) on the graph (Figure 6 right) we can additionally see that there is no evidence that these exotic species are interacting with natives, as they are estimated to be conditionally independent of them (Figure 6, middle), a point which was not clear from the negative correlations of Figure 6, left. We can also see that one of the exotic species, *Libertia* pulchella(LIBPUL) is more closely related to the native species. This may indicate that while the other exotic species are growing separate to native species, possibly in disturbed habitats, the *Libertia pulchella* is co-existing with native species, and may therefore be competing with natives for resources and space.

Curiously, we found that *Hymenophyllum sanguinolentum* (HYMSAN) and *Hymenophyllum villosum* (HYMVIL) have a negative partial correlation, even though they can cooccur (Brownsey & Perrie, 2014). These two species are often confused with each other. This negative relationship may therefore be an artifact of misclassification rather than a true negative relationship. Field ecologists may identify one or the other of these species, and assume everything similar in a plot is the same fern. The result would be that these species would rarely be recorded to co-occur.

We observe negative partial correlations between *Fuscospora cliffortioides* (NOTCLI) and other species when not controlling for any covariates (Figure 7a), which are lost after controlling for altitude and slope (Figure 7b). This is consistent with out understanding that *Fuscospora cliffortioides* occurs in montane and subalpine forest that tend to occur at altitudes between 400 m - 1380 m above sea level, whereas *Prumnopitys ferruginea* (PRUFER), *Weinmannia racemosa* (WEIRAC) and *Raukaua simplex* (RAUSIM) occur in lower altitude forest types, that range from sea-level up to 700 m in the south island and up to 1100 m in the north island (Wiser et al., 2011).

**Figure 7:**
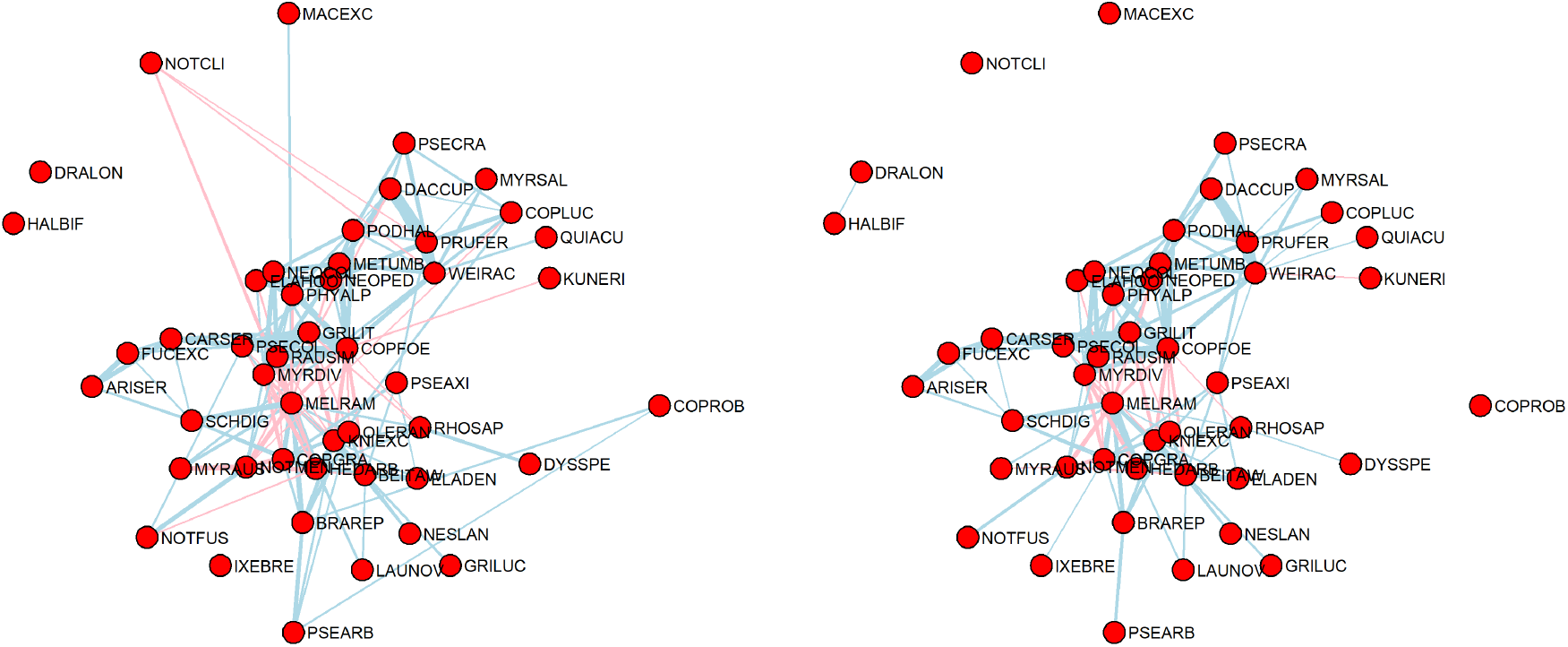
Partial correlation graph of trees before (left) and after (right) controlling for covariates. Looking at these graphs can suggest which correlations are explained by the covariates modelled. Negative partial correlations between *Fuscospora cliffortioides* (NOTCLI) and other species are not present after controlling for covariates (slope and altitude), presumably because *Fuscospora cliffortioides* tends to occur at higher altitudes.

## DISCUSSION

We have demonstrated a new method for exploring whether co-occurrence patterns between species may be explained by mediator species, or by response to environmental variables, or by neither. GCGMs can be used to analyze a wide variety of data types commonly observed in multi-species data in ecology, and can accommodate large datasets as well as small ones. We show, by simulation, that GCGMs are better able to distinguish effects of mediator species from direct associations than methods applied to transformed counts that assume normality. They are able reproduce known relationships between species, and can generate hypotheses based on surprising partial correlation patterns, as demonstrated in our analysis of the New Zealand native forest cover data.

Gaussian copulas, as used here, have exciting potential in ecology for the analysis of multivariate non-normal data. For example, the algorithm used to construct GCGMs here is quite general, and can be used to fit any desired covariance model on iteratively reweighted data (Popovic et al., 2018). Thus copulas can be readily used for ordination, using an iteratively reweighted factor analysis, as a fast, large-sample alternative to the hierarchical methods currently used for model-based ordination (Warton et al., 2015). Copulas can also be used as a simulation model (as in Warton et al., 2017), and have potential as a tool for likelihood-based inference about multivariate data. We look forward to seeing further progress using a copula approach for multivariate analysis in ecology.

## ACKNOWLEDGEMENTS

FKCH is supported by a Australia Research Council Discovery Project DP180100836 and a ANU cross-disciplinary research grant. DIW was supported by an Australian Research Council Future Fellowship (FT120100501). GCP was supported by the Australia Postgraduate Award and ARC Discovery Project scheme (DP180103543). ATM is supported by a Australia Research Council Discovery Grant (DP180100836).

Data was sourced from the New Zealand Ministry for the Environment Land Use Carbon Analysis System (LUCAS) Natural Forest Plot Measurement Programme.

## AUTHOR CONTRIBUTIONS

GCP carried out the data analysis and lead writing. DIW and FKCH provided ideas and input, and contributed critically to drafts. ATM and FJT assisted with interpretation of results from and ecological perspective and contributed to the writing for ecological aspects.

## DATA ACCESSIBILITY

Hunting spider data is available in the mvabund package in R. New Zealand native forest data is available from Manaaki Whenua Landcare Research (2018).

